# Long-term warming inverts the relationship between ecosystem function and microbial resource acquisition

**DOI:** 10.1101/2024.10.23.619826

**Authors:** Melissa S. Shinfuku, Kristen M. DeAngelis

## Abstract

Soil microbial traits drive ecosystem functions, which can explain the positive correlation between microbial functional diversity and ecosystem function. However, microbial adaptation to climate change related warming stress can shift microbial traits with direct implications for soil carbon cycling. Here, we investigated how long-term warming affects the relationship between microbial trait diversity and ecosystem function. Soils were sampled after 24 years of +5°C warming alongside unheated control soils from the Harvard Forest Long-Term Ecological Research site. Ecosystem function was estimated from six different enzyme activities and microbial biomass. Functional diversity was calculated from metatranscriptomics sequencing, where reads were assigned to yield, acquisition, or stress trait categories. We found that in organic horizon soils, warming decreased the richness of acquisition-related traits. In the mineral soils, we observed that heated soils exhibited a negative relationship with the richness of acquisition-related traits. These results suggest that microbial communities exposed to long-term warming are shifting away from a resource acquisition life history strategy.

---

Ecosystem function generally correlates with microbial diversity (Delgado-Baquerizo et al., 2020), and while taxonomic diversity is often used as a proxy for functional diversity, this underlying assumption is not supported for diverse communities, such as soils (Fierer et al., 2012). Functional diversity (FD) is usually a better predictor of ecosystem function than taxonomic diversity (Steudel et al., 2016), but measuring FD in microorganisms is challenging (Escalas et al., 2019). Function cannot be reliably inferred from taxonomy due to inconsistent phylogenetic conservation of traits (Martiny et al., 2013), horizontal gene transfer, and strain-level variation (Pan et al., 2023). To classify microbial traits into ecologically relevant life history strategies, the Yield, Acquisition, Stress (YAS) framework presents an adaptation of the Competitor, Stress tolerator, Ruderal framework (Malik et al., 2020). This traits-based approach to measuring FD better captures the trade-offs in microbial life history independent of taxonomy (Krause et al., 2014).

Chronic warming has depleted soil organic matter and altered microbial traits, so we sought to assess how long-term warming affects the relationship between functional diversity and ecosystem function. Since 1991, soils at the Harvard Forest warming experiment have been heated +5^*°*^C above ambient temperature (Melillo et al., 2017). Heating decreased total carbon (Pec et al., 2021) and specific compounds like lignin in the soil (Pec et al., 2021; Pold et al., 2017), possibly due to increased lignin decomposition (Pold et al., 2016). Hydrolytic enzyme abundance, a trait related to resource acquisition, is higher in the heated soils compared to controls (Anthony et al., 2021; Roy Chowdhury et al., 2021). Additionally, stress related traits such as biofilm formation may also increase with soil warming, since the heated soils are typically drier than the unheated control soils (Contosta et al., 2011). Though previously we found no significant effect of heating on the relationship between microbial taxonomic diversity and ecosystem function (Shinfuku et al., 2023), the underlying changes to substrate quality, substrate quantity, and microbial traits suggest that heating affects the relationship between ecosystem function and FD.

We hypothesized that long-term warming would have a positive effect on the relationship between functional diversity and ecosystem function, driven by increases in stress and resource acquisition associated traits. We also hypothesized that in the heated plots, stress-related traits would have a stronger relationship with EMF compared to the control plots.

Soils were collected on June 3^rd^ and October 27^th^ in 2014 from the Harvard Forest long-term soil warming experiment located in Petersham, MA after 24 years of warming, as previously described (Pold et al., 2017; Roy Chowdhury et al., 2021). The organic horizon and mineral soil in each soil core were separated in the field.

Ecosystem multifunctionality (EMF) was calculated from previously published extracellular enzyme activity (β-glucosidase, β-xylosidase, cellobiohydrolase, peroxidase, N-acetyl glucosaminidase, phenol oxidase) and microbial biomass data (Pold et al., 2017). EMF was calculated in the R package multifunc using the averaging approach described in Byrnes et al. (2014).

Functional diversity was calculated from previously published data (Roy Chowdhury et al., 2021). Raw reads were quality checked using FASTQC then merged using FLASh (Magoč and Salzberg, 2011). Merged reads were filtered using Trimmomatic (Bolger et al., 2014), and ribosomal RNA sequences removed using sortmerna (v.4.3.6) (Kopylova et al., 2012). Reads were aligned against the NCBI non-redundant (nr) protein database in DIAMOND (v.2.1.8) (Buchfink et al., 2021), and hits were translated with EMBOSS (v.6.6.0) (Rice et al., 2000) and assigned KEGG Orthology (KO) in MEGAN Ultimate Edition (v.6.25.5) (Huson et al., 2007). Reads were also queried in HMMER (v.3.3.2) against the CAZyme database, retrieved from dbCAN3 (Zheng et al., 2023). Reads were classified along the Yield Acquisition Stress (YAS) framework (Malik et al., 2020) based on KEGG pathway and CAZy annotation (Table 1). We calculated Shannon’s diversity, Chao1 estimated richness, and Pielou’s evenness for the yield, acquisition, and stress trait categories following a repeated rarefying approach (Schloss, 2024).

**Table 1:** KEGG pathways and CAZymes categorized into Yield, Acquisition, Stress framework.

| YAS category | KEGG pathway | Citation |
| --- | --- | --- |
| Yield | Basal transcription factors | Malik et al. (2020) |
|  | Ribosome | Li et al. (2022) |
|  | RNA polymerase | Krause et al. (2014); Piton et al. (2023) |
|  | Pyrimidine metabolism | Li et al. (2022); Ning et al. (2023) |
|  | Carbon fixation pathways in prokaryotes | Ning et al. (2023) |
|  | Pentose phosphate pathway | Malik et al. (2020); Ning et al. (2023) |
|  | Amino sugar and nucleotide sugar metabolism | Malik et al. (2020) |
|  | Aminoacyl-tRNA biosynthesis | Li et al. (2022); Ning et al. (2023) |
|  | Ribosome biogenesis in eukaryotes | Wood et al. (2018) |
|  | Glycolysis / Gluconeogenesis | Malik et al. (2020) |
|  | Citrate cycle (TCA cycle) | Malik et al. (2020) |
|  | Oxidative phosphorylation | Malik et al. (2020) |

Table 1: KEGG pathways and CAZymes categorized into Yield, Acquisition, Stress framework (Continued)
| YAS category | KEGG pathway | Citation |
| --- | --- | --- |
| Acquisition | Flagellar assembly | Malik et al. (2020); Ning et al. (2023); Piton et al. (2023) |
|  | Bacterial chemotaxis | Malik et al. (2020); Ning et al. (2023); Piton et al. (2023) |
|  | Biosynthesis of siderophore group nonribosomal peptides | Malik et al. (2020); Piton et al. (2023) |
|  | ABC transporters | Malik et al. (2020); Piton et al. (2023) |
|  | Glycoside hydrolases | Malik et al. (2020) |
| Stress | Exopolysaccharide biosynthesis | Malik et al. (2020) |
|  | Biofilm formation - *Vibrio cholerae* | Malik et al. (2020) |
|  | Biofilm formation - *Pseudomonas aeruginosa* | Malik et al. (2020) |
|  | Biofilm formation - *Escherichia coli* | Malik et al. (2020) |
|  | Mismatch repair | Li et al. (2022); Malik et al. (2020) |
|  | Homologous recombination | Li et al. (2022); Malik et al. (2020) |
|  | Base excision repair | Malik et al. (2020) |
|  | Two-component system | Malik and Bouskill (2022) |
|  | Glycine, serine, threonine metabolism | Malik et al. (2020) |

All statistical analyses were conducted in R (R Core Team, 2023). Diversity metrics, EMF, and linear model residuals were tested for normality using a Shapiro-Wilk test. We tested for differences in YAS trait diversity based on warming treatment or season using a Welch t-test. We constructed a set of linear models where EMF was the response variable and diversity, season, and warming were possible predictor variables. Models were assessed using ΔAICc, with the best fitting model determined by the lowest AICc score. Model errors are reported as a 95% confidence interval. Model p-values were Benjamini-Hochberg corrected to account for multiple comparisons. Adjusted p-values less than or equal to 0.05 were considered significant. Additional details for the methods can be found in the supplemental methods.

In heated organic horizon soils, the richness of acquisition traits was significantly lower compared to controls (Figure 1A). The decrease in acquisition related traits could be indicative of reduced carbon quality or quantity due to heating. For example, flagellar assembly is an acquisition trait that correlates with soil carbon availability (Ramoneda et al., 2024). Decreased acquisition trait richness could also indicate a shift towards a more oligotrophic lifestyle. Acquisition related traits are typically associated with fast-growing organisms that can respond quickly to increased substrate availability (Barnett et al., 2023; Piton et al., 2023). This shift in life history strategy reflects observed decreases in fungal biomass and diversity (Anthony et al., 2021;

**Figure 1.**
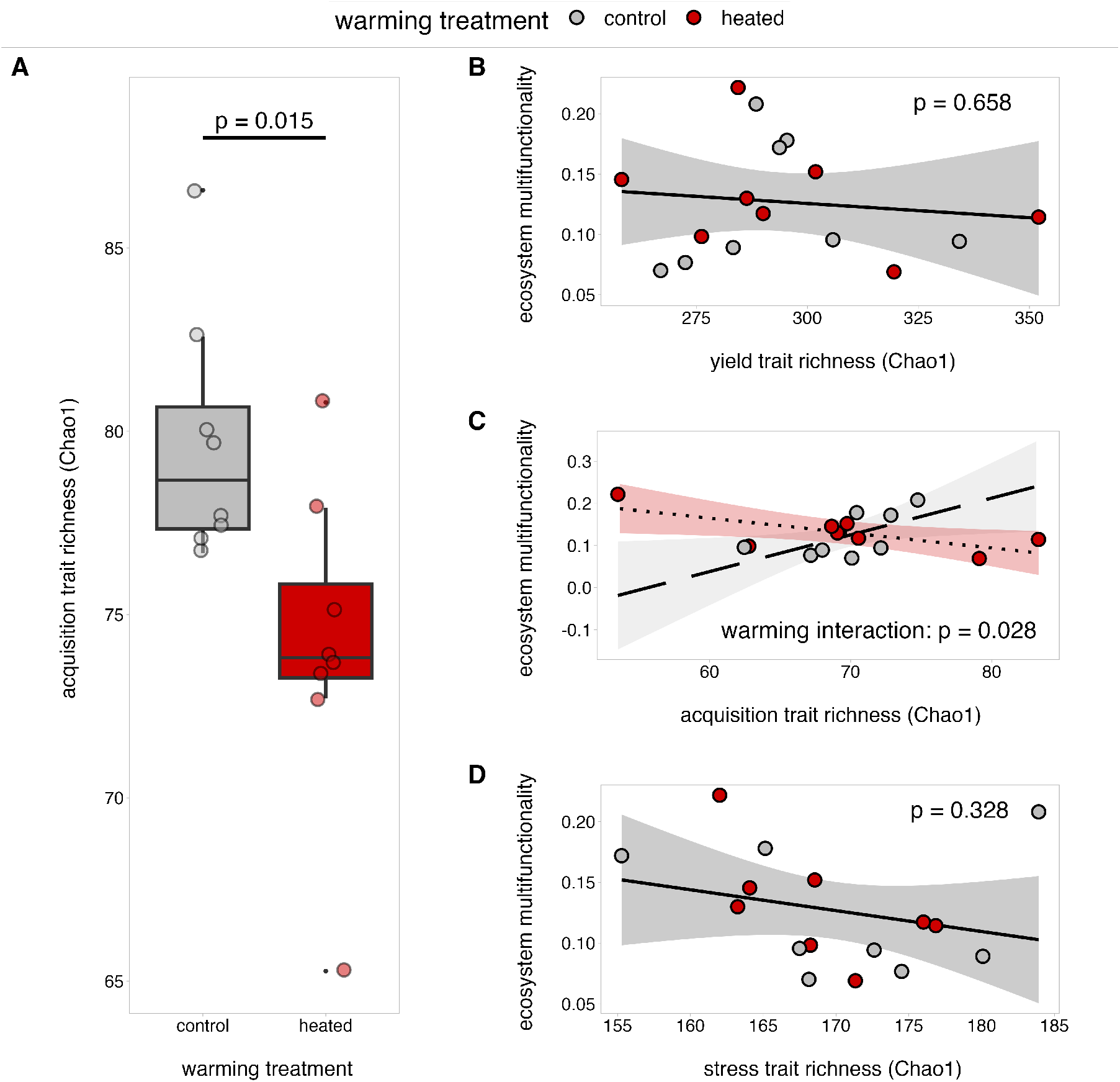
Yield, Acquisition, and Stress related traits in the organic horizon and mineral soils. Ecosystem multifunctionality (EMF) and acquisition trait richness were checked for normality using a Shapiro-Wilk test. Organic horizon acquisition trait richness (Chao1) in the heated and control soils was compared using a Welch’s t-test (A). Linear models were constructed to examined the relationship between EMF and yield (B), acquisition (C), and stress (D) trait richness. Model residuals were checked for normality using a Shapiro-Wilk test. Models were compared using ΔAICc. Any p-values for the linear model results were adjusted using a Benjamini-Hochberg correction, and the adjusted p-values are what is displayed on the figures. Shaded areas represent a 95% confidence interval.

Frey et al., 2008), along with a decrease in the average ribosomal RNA gene copy number in the heated plots (DeAngelis et al., 2015). There were no effects of heating treatment on the diversity of YAS traits in the mineral soils. Additionally, there were no significant differences in the diversity of the YAS traits between seasons in the organic or mineral soils.

In the mineral soils, the relationship between ecosystem multifunctionality and richness of acquisition associated traits had an interactive effect with warming (Figure 1C). Control soils showed a positive trend between EMF and richness of acquisition associated traits (0.009 ± 0.008, p = 0.062). Comparatively, in the heated plots, EMF and richness of acquisition associated traits had a significant negative relationship, contradictory to our hypothesis (−0.012 ± 0.008, p = 0.028). Trait richness for yield (Figure 1B) and stress (Figure 1D) had no significant relationship with EMF in mineral soils. In the organic horizon, none of the YAS trait category diversity metrics had a significant relationship with EMF either.

The negative relationship between acquisition trait richness and ecosystem function in the heated plots suggests that competition has a negative effect on ecosystem function. Acquisition traits are expressed to compete for resources (Malik et al., 2020), and competition can arise from lack of niche partitioning or complementarity (Cardinale, 2011), mechanisms which typically promote ecosystem function. Further, stress can reduce complementarity within a system (Baert et al., 2018). Higher levels of competition within a community can also produce negative selection effects, where traits unrelated to ecosystem function come to dominate a community because they provide a competitive advantage (Jiang et al., 2008).

In conclusion, warming had a negative effect on the richness of acquisition traits in the organic horizon, and resulted in a negative relationship between ecosystem function and acquisition traits richness in the mineral soils. Climate warming-related decreases in soil substrate quantity and quality may be shifting the microbial life strategy away from resource acquisition.

## Acknowledgements

This work was supported by a National Science Foundation (NSF) Long-Term Research in Environmental Biology grant (DEB-1456528), which funds the long-term warming experiment at the Harvard Forest. We are grateful to Drs. Serita Frey and Mel Knorr for maintaining this infrastructure for our community. This work was also supported by an NSF CAREER award (DEB-1749206) to KMD, and by the intramural research program of the U.S. Department of Agriculture, National Institute of Food and Agriculture, Hatch Multistate, accession number 7004345. The findings and conclusions in this publication have not been formally disseminated by the U. S. Department of Agriculture and should not be construed to represent any agency determination or policy.

