## Supplemental Methods for "Long-term warming inverts the relationship between ecosystem function and microbial resource acquisition"

### 1 Supplemental methods

#### 1.1 Site description and soil sampling

Since 1991, soils at the Harvard Forest long-term soil warming experiment have been continuously heated +5°C above ambient temperature using resistance cables buried 10 cm below the soil surface and spaced 20 cm apart (Peterjohn et al. 1994). The plant overstory is dominated by *Acer rubrum*, *Betula lenta*, *Betula papyrifera*, *Fagus grandifolia*, *Quercus velutina*, and *Quercus rubra*. The site receives an average of 1107 mm of precipitation, including snow, distributed evenly throughout the year (Boose et al. 2022). The mean annual temperature at the site ranges from a low of 3.3 °C to 13.1 °C (Boose et al. 2022). Both the organic horizon and mineral soils at the site are acidic and have pHs ranging from 3.8 to 4.2 and 3.9 to 4.4, respectively (Anthony et al. 2020; Pold et al. 2017). These pH values do not vary between the heated and control plots (Pec et al. 2021).

Duplicate or triplicate 10 cm cores were sampled from each plot using a 5.7 diameter tulip bulb corer, sterilized with 70% ethanol in between plots. The organic horizon and mineral soil were separated, replicates were pooled by soil depth, and roots and rocks were removed. The pooled soil samples were split into two parts; one part was flash frozen for lipid P analyses and the other part was transported back to the lab at ambient temperature for enzyme assays.

#### 1.2 Ecosystem function measurements

The potential enzyme activities of BG, BX, CBH, and NAG were assessed using a fluorescent assay with methylumbelliferyl substrates, and the potential enzyme activities of HPO and PO were assessed using a colorimetric assay. For the all of the enzyme assays, 1 g of fresh soil was combined with 125 ml of 50 mM sodium acetate buffer (pH 4.7) in a Waring blender to make a soil slurry. For the fluroescent enzyme assays, 200 µl of the soil slurry was combined with 50 µl of 100 µM of the methylumbelliferyl substrate in a black 96 well plate. Each plate also contained a standard curve, a substrate only control, and a soil only control. Plates were incubated at 25°C and read at 2, 4, and 6 hr after the addition of the substrate with excitation-emission set to 350/450 nm. For the colorimetric assays, equal parts of the soil slurry and 25 mM substrate solution were combined (L-DOPE for phenol oxidase and L-DOPE + 0.3% hydrogen peroxide for peroxidase).

For measuring microbial biomass via total lipid phosphate, 0.3 g of organic soil or 1 g of mineral soil were extracted in a 2:1:0.8 mixture of methanol:chloroform:citrate buffer (0.15 M pH 4) in duplicate (Findlay et al. 1989). After a 12 hr extraction, lipids were digested using a saturated postassium persulfate solution at 95°C for 36 hr, with β-glycerophosphate digested additionally as a standard.

For the averaging approach, ecosystem function measurements were z-score transformed, then the measures were averaged for each sample. This yielded an ecosystem multifunctionality index for each sample.

#### 1.3 Sequence processing

After quality checking in FASTQC, forward and reverse reads were then merged using FLASh (Magoč et al. 2011), setting the maximum overlap parameter (-M) to 150. Reads were trimmed in Trimmomatic (Bolger et al. 2014), with the following parameter settings: TRAILING:33 and LEADING:33. For removing ribosomal RNA sequences, we used SILVA 138 SSURef NR99, SILVA 138 LSURef, and 5S and 5.8S RFAM databases. In total, about 91% of the trimmed reads were identified as ribosomal RNA sequences and removed. The NCBI non-redundant protein database was retrieved on July $7^{th}$, 2023. Of the 295,487,595 non-ribosomal RNA reads, 77,956,651 reads aligned with a protein sequence in the NCBI non-redundant protein database. To translate the putative mRNA sequences into reading frames, we used function 'transeq' in EMBOSS (v 6.6.0) (Rice et al. 2000). Only reads that had e-values $<$ 1e-3 and coverage fractions $>$ 0.3 were retained. If multiple reading frames from a single sequencing read aligned with different regions within a CAZyme sequence, the reading frame with the lowest e-value was selected. Any CAZymes that appeared once across all samples were discarded. After querying the reading frames against the CAZyme database, only CAZymes in the auxilary activities (AA), carbohydrate binding modules (CBM), carbohydrate esterases (CE), glycoside hydrolases (GH), glycosyltransferases (GT), or polysaccharide lyases (PL) categories were retained, which summed to 127,648 CAZymes. In total, 28,591,960 out of 77,956,651 putative mRNA reads were matched with a KEGG Ortholog. Any KOs which were observed once across all of the samples were removed (1349 KOs in total). The final KO count was 28,590,611.

KEGG Orthologs were also assigned to either yield, acquisition, or stress based on involvement in specific KEGG pathways, which were pulled from the KEGG database on June 7, 2024 using the R package KEGGREST (Tenenbaum et al. 2022). In the KEGG database, KOs have been assigned or associated with a specific or multiple pathways. These pathways can contain single or multiple KOs. Using the KEGG pathway assignments, we then investigated KOs that were assigned to one of the pathways of interest. Of the 17173 unique KOs in the final dataset, 1351 belonged to at least one of the YAS pathways, and of the 361 unique CAZys, 135 were identified as glycoside hydrolases.

#### 1.4 Functional diversity calculations

To assess functional diversity, we calculated Shannon's diversity, Chao1 estimated richness, and Pielou's evenness diversity metrics for the assigned transcripts based on the counts of individual KEGG Orthology numbers. Prior to calculating the diversity metrics, we rarefied samples to 7493 KO counts, then calculated diversity metrics. This process was repeated 1000 times and the mean of those diversity metrics is what was utilized for further analysis. While the validity of rarefaction has been questioned (McMurdie et al. 2014), recent papers have found it to be the best way to address sequencing depth disparities between samples and rarefying multiple times tends to perform better than rarefying once when effect sizes are small (Schloss 2024). Since previous work observed minor effects of warming on microbial diversity (DeAngelis et al. 2015; Shinfuku et al. 2023), we elected to rarefy multiple times as a way to account for uneven sequencing depth across the samples. Since we wanted to compare the yield, acquisition, and stress diversity metrics, we rarefied to the lowest counts observed across all three categories, which was 229 counts.

##### 1.5 Statistical analyses

For generalized linear model construction, in both the organic horizon and mineral soils, EMF was normally distributed, so all models utilized a Gaussian distribution. We considered the following predictor combinations: diversity, warming + diversity, season + diversity, warming + diversity, warming + season + diversity, and all possible interactions between these variables. We elected to always include diversity as a predictor since we are interested in examining the relationship between KEGG, CAZyme, or YAS trait diversity and EMF.

When comparing models, the best fitting model was determined as the model with the lowest AICc score that was at least 2 ΔAICc units greater than the next best scoring model. If the best scoring models were less than 2 ΔAICc units apart, a likelihood ratio test was used to compare the models if they were nested. If the similar scoring models were not nested, then model with the lowest residual deviance was selected.
